# Activity deprivation modulates the Shank3/Homer1/mGluR5 signaling pathway to enable synaptic upscaling

**DOI:** 10.1101/2025.04.24.650518

**Authors:** Andrea A. Guerrero, Gina G. Turrigiano

**Affiliations:** Brandeis University

## Abstract

Shank3 is an autism spectrum disorder-associated postsynaptic scaffold protein that links glutamate receptors to trafficking and signaling networks within the postsynaptic density. Shank3 is required for synaptic scaling (Tatavarty et al., 2020), a form of homeostatic plasticity that bidirectionally modulates post-synaptic strength in the right direction to stabilize neuronal activity. Shank3 undergoes activity-dependent phosphorylation/dephosphorylation at S1586/S1615, and dephosphorylation at these sites is critical for enabling synaptic upscaling (Wu et al., 2022). Here, we probe the molecular machinery downstream of Shank3 dephosphorylation that allows for synaptic upscaling. We first show that a phosphomimetic mutant of Shank3 has reduced binding ability and interaction with long-form Homer1, a postsynaptic protein also crucial for scaling, and a known binding partner of Shank3. Since metabotropic glutamate receptor 5 (mGluR5) has been shown to associate with Shank3 through long-form Homer1, we manipulated mGluR5 signaling with either noncompetitive or competitive inhibitors and found that only competitive inhibition (which targets agonist-dependent signaling) impairs synaptic upscaling. Further, we found that mGluR5 activation rescues synaptic upscaling in the presence of phosphomimetic Shank3, thus is downstream of Shank3 phosphorylation. Finally, we identify necessary signaling pathways downstream of group I mGluR. Taken together, these data show that activity-dependent dephosphorylation of Shank3 remodels the Shank3/Homer1/mGluR signaling pathway to favor agonist-dependent mGluR signaling, which is necessary to enable synaptic upscaling. More broadly, because downscaling depends on agonist-*independent* mGluR5 signaling, these findings demonstrate that synaptic up and downscaling rely on distinct functional configurations of the same signaling elements.

**SIGNIFICANCE STATEMENT:** Synaptic scaling is a bidirectional, homeostatic form of synaptic plasticity that allows neural circuits to maintain stable function in the face of experience-dependent or developmental perturbations. Synaptic scaling up requires dephosphorylation of the Autism Spectrum Disorder (ASD)-associated synaptic scaffold protein Shank3, but how this dephosphorylation event enables scaling up was unknown. Here we show that dephosphorylation of Shank3 rearranges interactions between synaptic proteins to drive agonist-dependent signaling through metabotropic glutamate receptors (mGluRs), and that this signaling is necessary for scaling up. These findings show that altered mGluR signaling is downstream of Shank3 during homeostatic plasticity, and raise the possibility that some human Shankopathies impair signaling through this important signaling pathway.

## INTRODUCTION

Synaptic scaling is a form of homeostatic plasticity that bidirectionally adjusts synaptic weights in response to prolonged changes in firing, in the right direction to stabilize network activity (Turrigiano et al., 1998; Turrigiano and Nelson, 2004; Turrigiano, 2008). A large body of work has established that synaptic scaling is expressed through changes in the postsynaptic accumulation of AMPA-type glutamate receptors (AMPAR, O’Brien et al., 1998; Gainey et al., 2009, 2015; Goold and Nicoll, 2010; Louros et al., 2018) and is accompanied by a complex remodeling of postsynaptic scaffolds (Hu et al., 2010; Sun and Turrigiano, 2011; Steinmetz et al., 2016; Venkatesan et al., 2020), but the pathways that link changes in activity to changes in synaptic AMPAR accumulation are incompletely understood. We recently showed that activity blockade reduces Shank3 phosphorylation at two key sites (S1586 and S1615), and this dephosphorylation is necessary for the expression of synaptic scaling up (Wu et al., 2022). Here, we focus on identifying the molecular pathways downstream of Shank3 phosphorylation that enable synapses to undergo upscaling. We find that the phosphorylation state of Shank3 regulates binding to long-form Homer1, an interaction partner that links Shank3 to metabotropic glutamate receptors (mGluRs, Tu et al., 1999; Hayashi et al., 2009), and that agonist-dependent mGluR5 signaling is an essential downstream mediator of scaling up.

Shank3 is a multidomain scaffolding protein found abundantly at the postsynaptic density (Naisbitt et al., 1999), that interacts with several scaffold and signaling proteins known to be crucial for synaptic scaling (Grabrucker et al., 2011; Jiang and Ehlers, 2013). Synaptic upscaling depends on the dephosphorylation of two Serine sites (rat S1586/S1615) within Shank3, while synaptic downscaling requires phosphorylation of these same sites (Wu et al., 2022). One notable Shank3 interaction partner is Homer1, which has short and long isoforms - short-form Homer1a and long-form Homer1b/c. While long-form Homer1 is able to crosslink various proteins including Shanks and mGluRs, Homer1a interrupts this binding and promotes constitutive agonist-independent mGluR activity (Brakeman et al., 1997; Tu et al., 1999; Ango et al., 2001). Enhanced network activity leads to an increase in Homer1a that then shifts the balance of long-form to short-form Homer1 (Hu et al., 2010; Diering et al., 2017). This results in increased Homer1a-mGluR5 interactions (Lautz et al., 2018; Stillman et al., 2022) that lead to mGluR agonist-independent activity, thus enabling synaptic downscaling (Hu et al., 2010; Diering et al., 2017). In contrast, synaptic upscaling results in increased proteomic network associations between Shank3 and long-form Homer1, and between long-form Homer1 and mGluR5 (Heavner et al., 2021). Therefore, we hypothesized that Shank3 dephosphorylation facilitates synaptic upscaling by modulating its interaction with long-form Homer1, thus altering signaling through group I mGluR.

To test this hypothesis, we use quantitative immunohistochemical, biochemical, and pharmacological approaches to assess the role of long-form Homer1 and group I mGluR signaling in synaptic scaling up. We found that scaling up protocols produced correlated increases in pre- and postsynaptic proteins, and enhanced colocalization between endogenous Shank3 and long-form Homer1. This enhanced colocalization was prevented by overexpressing a phosphomimetic form of Shank3 that has reduced binding to long-form Homer1, suggesting that dephosphorylation during activity-blockade is necessary for this process. We then used pharmacology to show that - in contrast to downscaling - agonist-*independent* activity of group I mGluRs was not necessary for upscaling; instead, agonist-*dependent* activity was the critical factor. Further, enhancing mGluR5 signaling with a positive allosteric modulator was able to rescue synaptic upscaling in neurons overexpressing phosphomimetic Shank3, indicating that this signaling is downstream of Shank3 phosphorylation. Finally, we demonstrated that phospholipase C (PLC) and protein kinase C (PKC), downstream effectors of the canonical group I mGluR pathway, are required for synaptic scaling up. Taken together, these data show that activity-dependent dephosphorylation of Shank3 remodels the Shank3/Homer1/mGluR signaling pathway to favor agonist-dependent mGluR signaling, and this is necessary for the induction of synaptic scaling up.

## MATERIALS AND METHODS

### Neuronal cultures, transfection, and drug treatments

Primary neuronal cultures were prepared from the visual cortex of Long-Evans rat pups (postnatal days 1–3) from timed-pregnant Dams from Charles River. Neurons were dissociated from primary visual cortex and plated onto glass-bottomed dishes pre-seeded with glial feeders, following previously described methods (Tatavarty et al., 2020; Wu et al., 2022). All experiments were performed 7 to 10 days in vitro (DIV). During this period, neurons were sparsely transfected with the following constructs using Lipofectamine 2000 (Thermo Fisher Scientific). To express Shank3 phospho-mutants exogenously, GFP-tagged Shank3 constructs (2500 ng per dish) were transfected . For better visualization of neurons during imaging, an empty GFP vector was co-transfected in both conditions. In imaging experiments measuring endogenous Shank3 and Homer1b/c, an empty GFP vector (500 ng per dish) was transfected to delineate the neurons.

For each pharmacology experiment, sister cultures were treated with vehicle as a control. Neurons aged DIV 7-8 were treated with 4µM tetrodotoxin (TTX) to induce synaptic scaling up *in vitro* (Turrigiano et al., 1998). The drug/vehicle was added to the cultures at the same time as tetrodotoxin (TTX) for a total of 24 or 30 hours before fixation.

In all imaging experiments, we analyzed only excitatory pyramidal neurons, identified by their stereotypical morphology (Pratt et al., 2003). All experiments were replicated a minimum of three times from independent dissociations, with the exception of the MTEP data which are from two dissociations.

### Expression constructs

The construct expressing wild-type rat Shank3 (AJ133120.1) with an N-terminal GFP tag was obtained from Chiara Verpelli (Verpelli et al., 2011). Constructs expressing GFP-tagged Shank3 phospho-mutants were generated using the Gibson Assembly kit (NEB) with the wild-type Shank3 as the template (Wu et al., 2022). The same process was utilized to generate the GFP-tagged Shank3 P1311L point mutant and HA-tagged Homer1c, but the latter was derived from Addgene construct 54912 as a template. All constructs were verified by DNA sequencing.

### HEK cell cultures and transfection

HEK 293T cells (CRL-3216, ATCC) were plated from frozen stocks onto 10 cm plates and grown in media consisting of 84% DMEM, 10% FBS, 5% Pen-Strep, and 1% L-Glutamine. The HEK cells were passaged 3 times before plating onto 6 well plates. Transfection with Lipofectamine 2000 was performed when approximately 85% confluence was reached. HEK cells were co-transfected with GFP empty vector (500ng per well) or a GFP-tagged Shank3 construct (2450ng per well) and wildtype HA-tagged Homer1c construct (1450ng per well).

### Co-immunoprecipitation

At 24 hours post-transfection, the HEK cells were washed three times with PBS (Gibco) and lysed with RIPA buffer containing 150mM NaCl, 50mM TrisHCl, 1mM EDTA, 1% Triton X-100, 0.1% SDS, 0.5% sodium deoxycholate, 1 tablet of protease inhibitor cocktail (Roche), and 1 tablet of PhosSTOP phosphatase inhibitor cocktail (Roche). After sonication and centrifugation, lysates were incubated either with goat-anti-rabbit GFP (A-11122) overnight followed by a 2 hour incubation with Protein G Dynabeads (ThermoFisher 1003D) or GFP-trap agarose beads or GFP-trap magnetic particles (Chromtek gtma-20, M-270) for 2 hours at 4°C. The beads/particles were washed with RIPA buffer three times before eluting in 30uL of 1x LD Buffer (Invitrogen LC2570) at 95°C for 10 minutes. Lysates were run on 7% Tris-Acetate gels (Invitrogen EA0358) and then transferred overnight at 4°C. Blots were blocked using Intercept PBS Blocking buffer (Licor) for 1 hour at room temperature and then incubated with guinea pig Shank3 (Synaptic Systems 162 304) and Anti-HA-Biotin (Roche 12158167001), or goat-anti-rabbit GFP (A-11122) and mouse Homer1 (Synaptic Systems 160 011) antibodies. The next day, blots were washed three times with PBS-T and incubated for 1 hour with guinea pig 680RD and streptavidin 800CW, or rabbit 680RD and mouse 800CW IR dye secondary antibodies (Jackson Labs). Western blot images were taken using BioRad imager and quantified using ImageJ (NIH).

### Immunocytochemistry

Forty-eight hours post-transfection, neurons were fixed with 4% paraformaldehyde for 15 minutes and processed for immunocytochemistry using an established protocol (Tatavarty et al., 2020; Wu et al., 2022). To determine the synaptic localization of endogenous or exogenously expressed Shank3, cells were permeabilized with blocking buffer (0.1% Triton X-100/5% goat serum in PBS) at room temperature for 30 minutes. They were then incubated overnight at 4°C in dilution buffer (5% goat serum in PBS) containing the following primary antibodies: chicken anti-GFP (1:1000, Aves Labs GFP 10-20), guinea pig anti-Shank3 (1:50, Cell Signaling #64555), and guinea pig Homer1b/c (1:500, Synaptic Systems 160 025). To visualize the surface GluA2, the protocol was modified. Prior to permeabilization, cells were first incubated for 90 minutes at room temperature with mouse anti-GluA2 (1:1000, gift from Eric Gouaux, Vollum Institute, Oregon Health & Science University, Portland, OR) diluted in blocking buffer without Triton X-100. Then, cells were permeabilized and incubated overnight at 4°C in dilution buffer (5% goat serum in PBS) with chicken anti-GFP (1:1000, Aves Labs GFP 10-20) and anti-VGluT1 (1:1000, Synaptic Systems 135 304).

The next day, neurons were washed three times with PBS and incubated for 1 hour at room temperature with dilution buffer containing the following secondary antibodies (1:400, Thermo Fisher Scientific): goat anti-chicken Alexa-488, goat anti-mouse Alexa-555, goat anti-guinea pig Alexa-555, goat anti-guinea pig Alexa-647, and goat anti-rabbit Alexa-647. After three more PBS washes, the glass bottoms containing stained neurons were detached from the dishes, mounted onto slides using DAPI-Fluoromount-G mounting medium (SouthernBiotech), and sealed with nail polish.

### Image acquisition and analysis

All images were acquired using a Plan-Apochromat 63×/1.40 oil objectives or a laser-scanning Fast AiryScan confocal microscope (LSM 880 FAS, Zeiss), with the exception of images in Figures 5C-F that were acquired on laser-scanning confocal microscope (LSM 880, Zeiss), using ZEN Black acquisition software. Acquisition settings—including laser power, gain/offset, and pinhole size—were kept consistent across all experiments. During image acquisition, pyramidal neurons were identified by their characteristic teardrop-shaped somata and apical-like dendrites.

For each neuron, approximately 15 stacked images (step size: 0.18 µm) were obtained to capture the apical dendrites and their branches, then subjected to maximum intensity projection using ZEN Black. To quantify synaptic protein colocalization and signal intensities, images were analyzed using MetaMorph software (Molecular Devices). A region of interest was manually drawn to include dendritic branches distal to the primary branch point of the apical-like dendrite. Signal intensity thresholds were set for each channel to exclude background noise, maintaining consistency across experimental groups. MetaMorph’s granularity function was used to threshold puncta in each channel (puncta size: 0.5–5 µm). Binary images were generated to outline identified puncta, and colocalized puncta were determined using the Logical AND operation.

A synapse was defined as a punctum double-labeled with sGluA2 and VGluT1 (Figures 1, 3G, 5, and 6) or triple-labeled with GFP, sGluA2, and VGluT1 (Figure 3G). Total puncta intensity for each channel at colocalized sites was then measured. Colocalization between total Shank3 and Homer1b/c was quantified for GFP-transfected neurons (Figure 2). GFP-labeled Shank3 and Homer1b/c was quantified for neurons overexpressing Shank3 DD (Figure 4).

**Figure 1.**
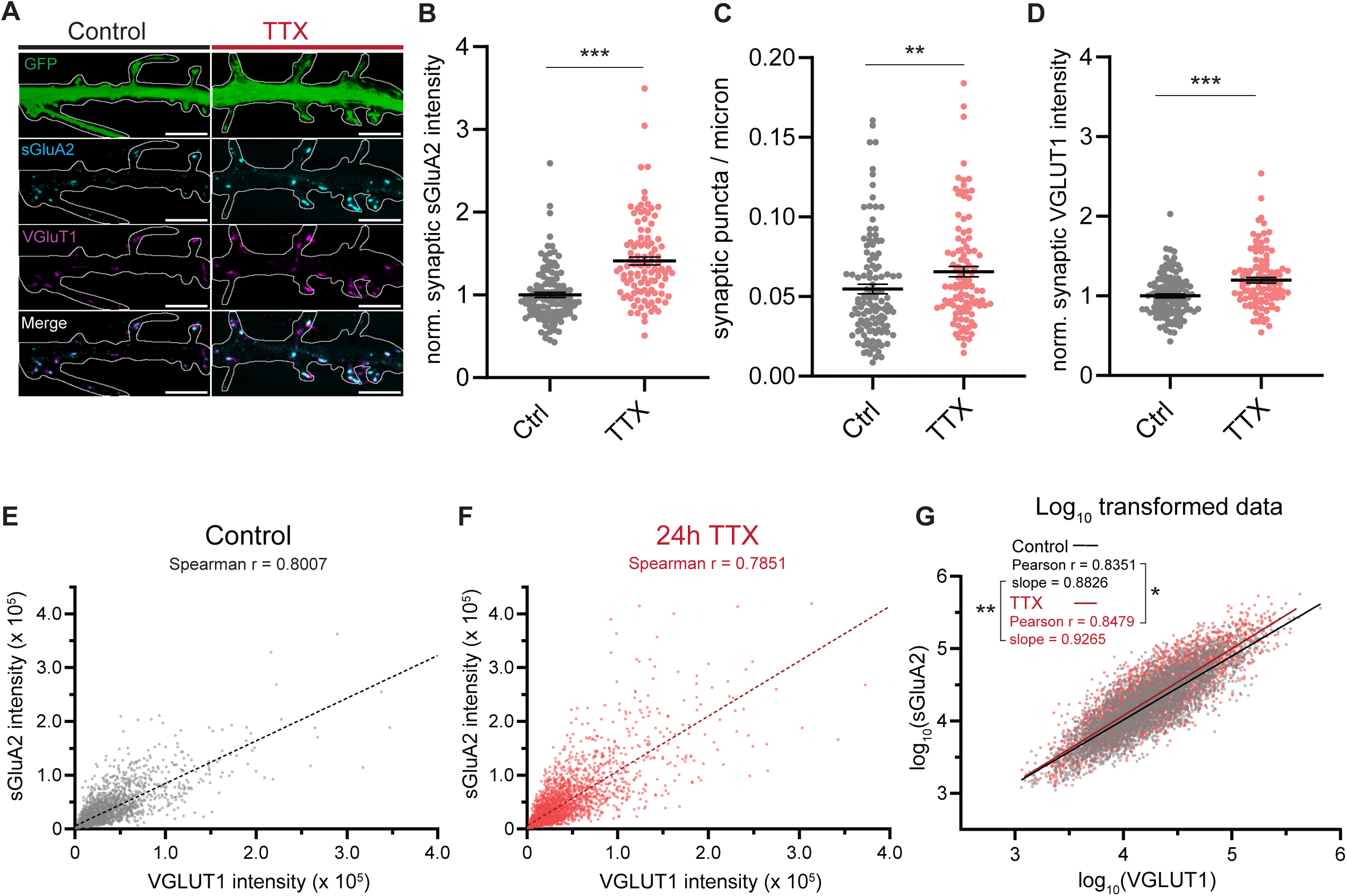
Synaptic scaling up protocols enhance excitatory synaptic density and produce correlated changes in pre- and post-synaptic proteins. ***A***, Representative images of sGluA2 and VGluT1 puncta in neuron expressing GFP and treated with DMSO vehicle (Ctrl) ± tetrodotoxin (TTX) (scale bar = 5 µm). ***B***, Quantification of synaptic sGluA2 intensity changes induced by scaling up protocol (number of neurons: control, n=120, TTX, n=104, Mann-Whitney U test: Ctrl vs. TTX, ***p<0.0001). Solid horizontal black lines indicate mean, and error bars represent SEM. ***C***, Quantification of putative synaptic puncta density changes induced by scaling up protocol (number of neurons: control, n=120, TTX, n=104, Mann-Whitney U test: Ctrl vs. TTX, **p=0.0036). ***D***, Quantification of synaptic VGLUT1 intensity changes induced by scaling up protocol (number of neurons: control, n=120, TTX, n=104, Mann-Whitney U test: Ctrl vs. TTX, ***p<0.0001). ***E***, VGLUT1 intensity plotted against sGluA2 intensity for control condition. Linear best fit of non-normally distributed raw data shown in dotted line (number of puncta: UN, n=4398). ***F***, VGLUT1 intensity plotted against sGluA2 intensity for 24 hour TTX treatment condition. Linear best fit of non-normally distributed raw data shown in dotted line (number of puncta: TTX, n=4658). ***G***, Log_10_ transformed data from E and F. Line of best-fit for each condition shown in solid lines (F test to determine lines are significantly different, line of best fit slope, **p=0.0003). Fisher’s z-transformation was used to compare Pearson’s correlation coefficients (*p=0.0457).

**Figure 2.**
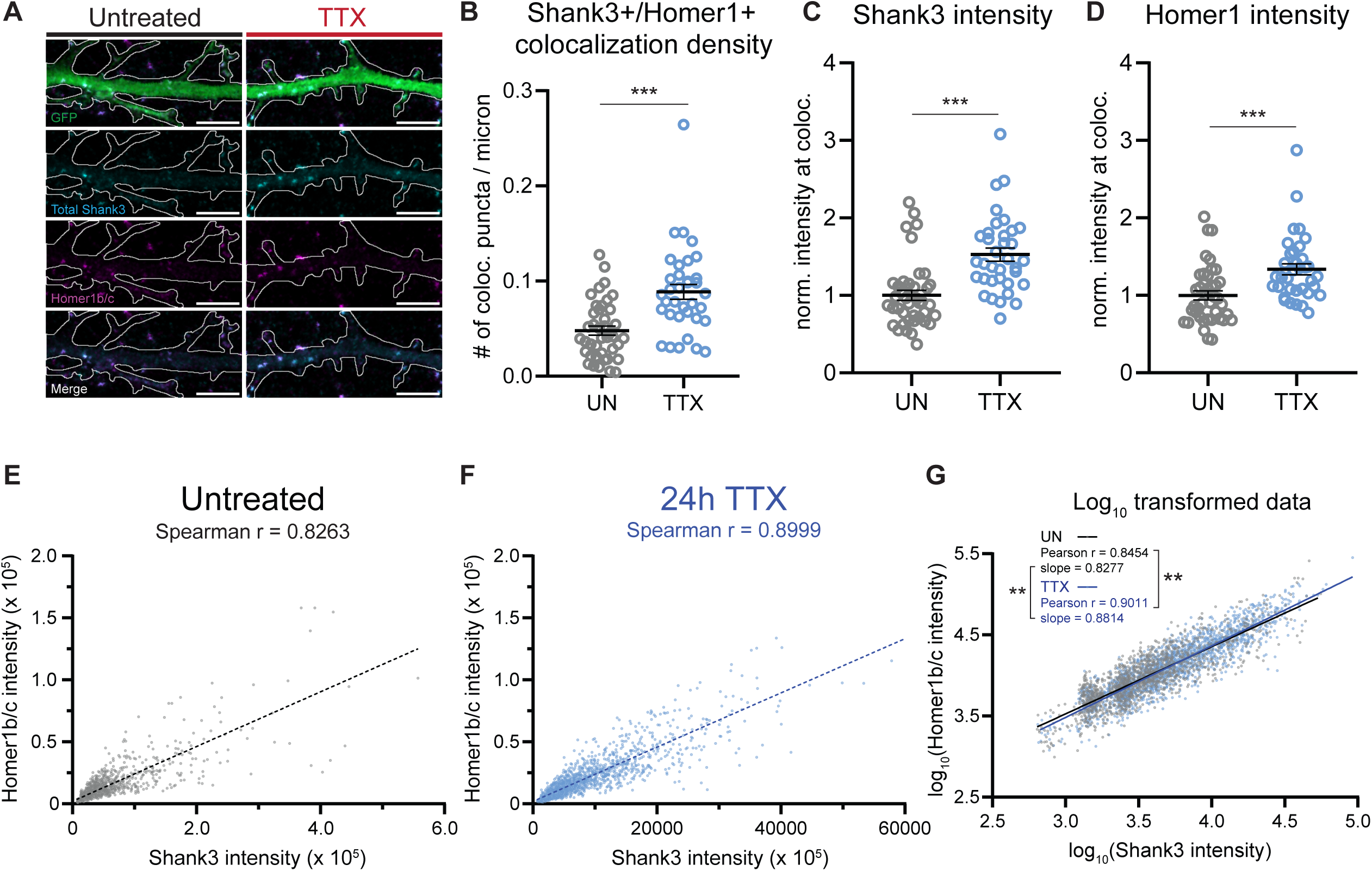
Synaptic scaling up protocols enhance endogenous Shank3-Homer1 colocalization. ***A***, Representative images of Shank3 and Homer1b/c puncta in dendrites from neurons expressing GFP (UN) ± tetrodotoxin (TTX) (scale bar = 5 µm). ***B***, Quantification of changes in Shank3+ Homer1b/c+ puncta density induced by scaling up protocol (number of neurons: untreated, n=41, TTX, n=35, Mann-Whitney U test: UN vs. TTX, ***p<0.0001). ***C***, Quantification of synaptic Shank3 intensity changes induced by scaling up protocol (number of neurons: untreated, n=41, TTX, n=35, Mann-Whitney U test: UN vs. TTX, ***p<0.0001). ***D***, Quantification of synaptic Homer1b/c intensity changes induced by scaling up protocol (number of neurons: untreated, n=41, TTX, n=35, Mann-Whitney U test : UN vs. TTX, ***p=0.0002). ***E***, Shank3 intensity plotted against Homer1 intensity for untreated condition. Linear best fit of non-normally distributed raw data shown in dotted line (number of puncta: UN, n=1511). ***F***, Shank3 intensity plotted against Homer1 intensity for 24 hour TTX treatment condition. Linear best fit of non-normally distributed raw data shown in dotted line (number of puncta: TTX, n=2228). ***G***, Log_10_ transformed data from E and F. Line of best-fit for each condition shown in solid lines (F test to determine lines are significantly different, line of best fit slope, **p=0.0006). Fisher’s z-transformation was used to compare Pearson’s correlation coefficients (***p<0.0001).

**Figure 3.**
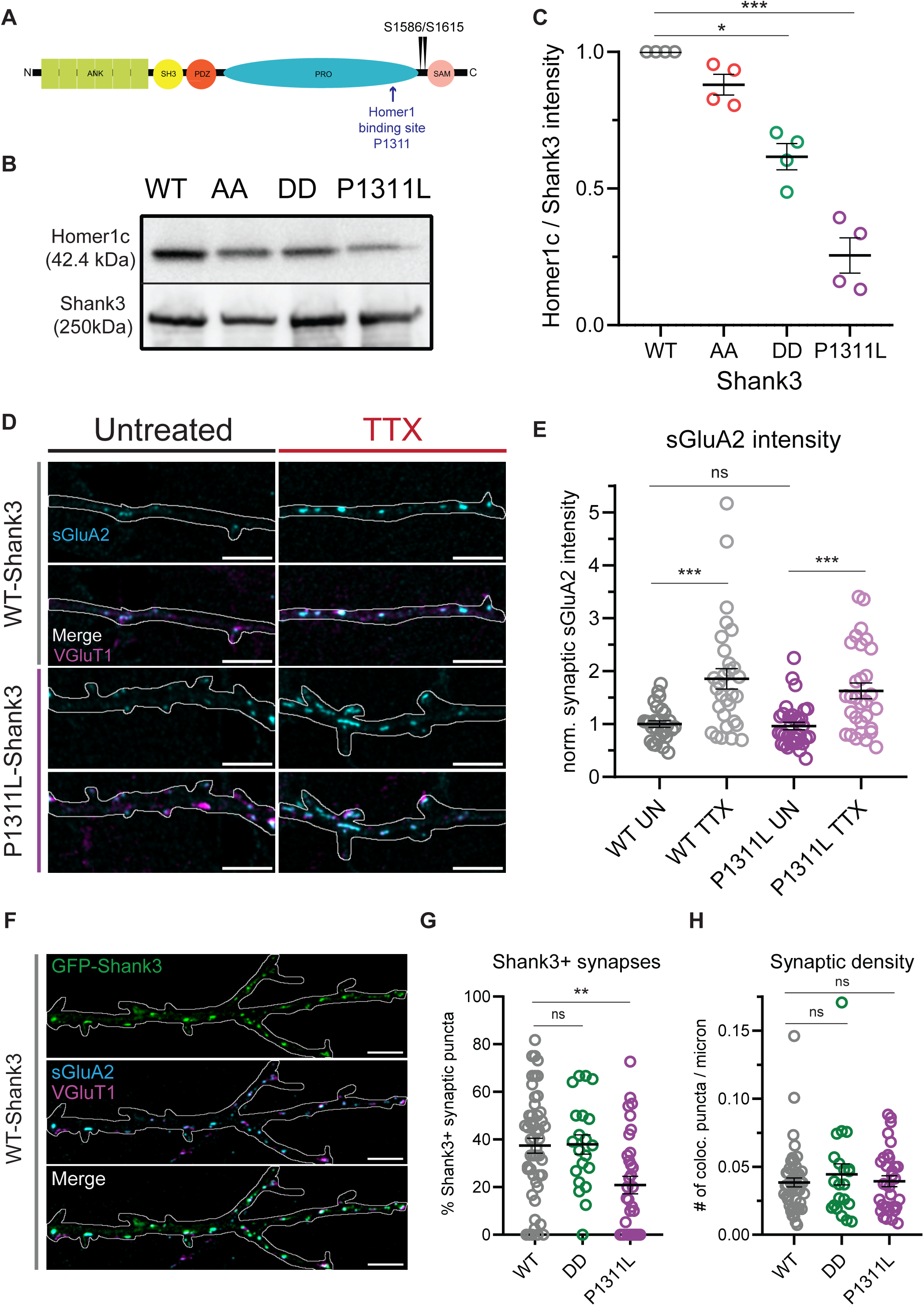
Phosphomimetic Shank3 DD has reduced interaction with Homer1b/c but maintains its ability to localize to the synapse. ***A***, Schematic of full-length Shank3 (Shank3 WT) with various domains, Shank3 phosphorylation sites of interest, and P1311L point mutation location labeled. ***B***, Representative Western blot image of Homer1c and Shank3 bands corresponding to Shank3 pulldowns of lysate from HEK cells overexpressing various Shank3 constructs. ***C***, Quantification of Western blot from Shank3 pulldowns by plotting Homer1c intensity value divided by corresponding Shank3 band intensity (4 biological replicates with each normalized to Homer1c band intensity divided by WT Shank3 band intensity value, Kruskal–Wallis test with post-hoc Dunn’s multiple comparisons test: WT vs. AA, ns p=0.6939, WT vs. DD, *p=0.05, WT vs. P1311L, **p=0.001). ***D***, Representative images of GFP-tagged Shank3, VGluT1, and sGluA2 puncta from WT and P1311L Shank3 overexpressing neurons. ***E***, Quantification of synaptic sGluA2 intensity changes induced by scaling up protocol in WT and P1311L Shank3 overexpressing neurons (UN) ± tetrodotoxin (TTX) (number of neurons: WT UN, n=28, WT TTX, n=30, P1311L UN, n=33, P1311L TTX, n=30, Kruskal–Wallis test with post-hoc Dunn’s multiple comparisons test: WT UN vs. P1311L UN, ns p>0.9999, WT UN vs. WT TTX, ***p=0.0005, P1311L UN vs. P1311L TTX, ns ***p=0.0005). ***F***, Representative images of GFP-tagged Shank3, VGluT1, and sGluA2 puncta from WT, DD, and P1311L Shank3 overexpressing neurons. ***G***, Quantification of percentage Shank3+ synapses out of total number of synapses represented by sGluA2 and VGluT1 colocalizations in WT, DD, and P1311L Shank3 overexpressing neurons (number of neurons: WT, n=50, DD, n=21, P1311L, n=33, Kruskal– Wallis test with post-hoc Dunn’s multiple comparisons test: WT vs. DD, ns p>0.9999, WT vs. P1311L, **p=0.0017). ***H***, Quantification of synaptic density in WT, DD, and P1311L Shank3 overexpressing neurons (number of neurons: WT, n=50, DD, n=21, P1311L, n=33, Kruskal–Wallis test with post-hoc Dunn’s multiple comparisons test: WT vs. DD, ns p>0.9999, WT vs. P1311L, ns p>0.9999).

**Figure 4.**
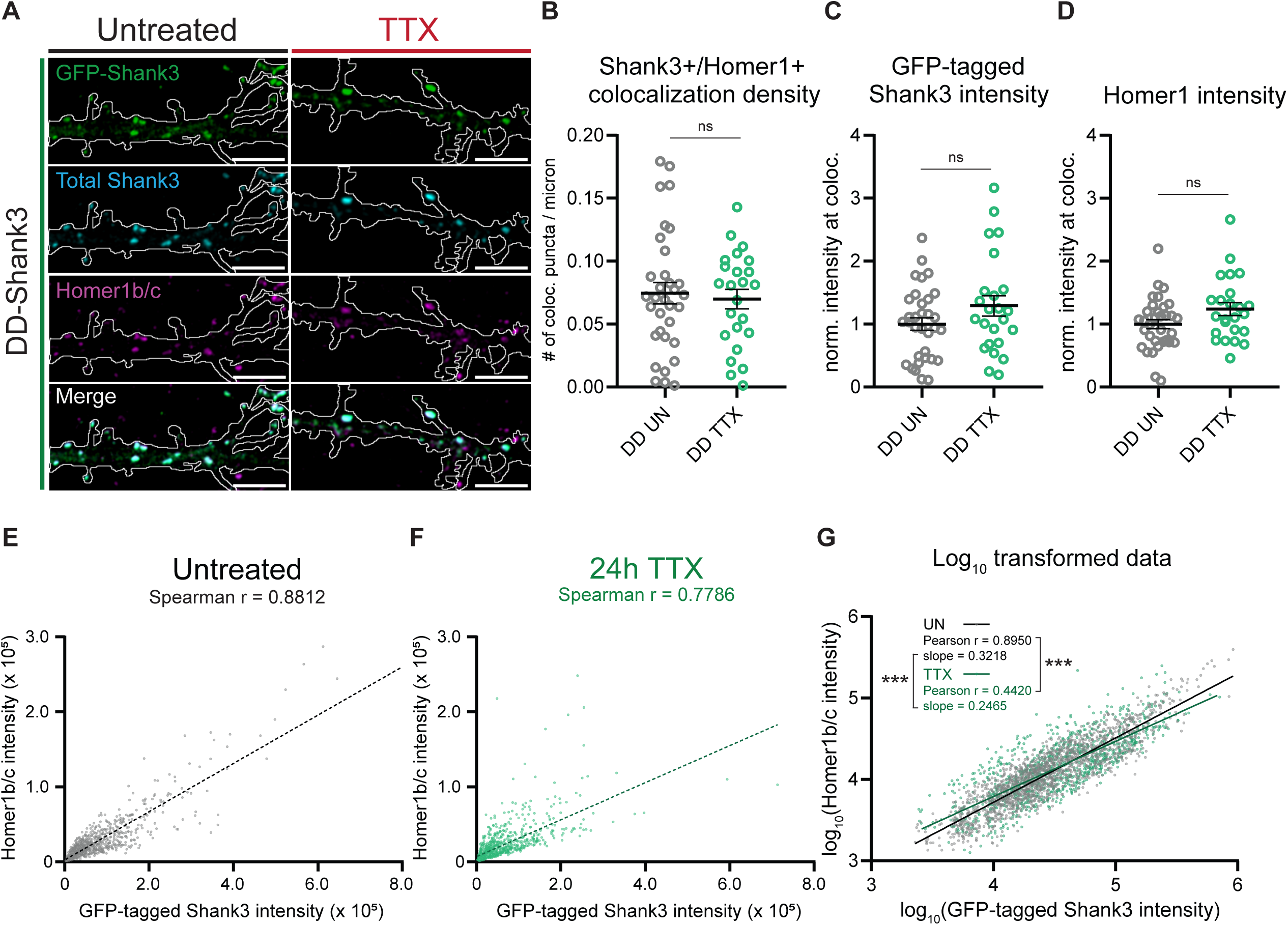
Synaptic scaling up protocols do not alter Shank3-Homer1 colocalization nor produce correlated changes between these proteins in the presence of phosphomimetic Shank3 DD. ***A***, Representative images of GFP-tagged Shank3 and Homer1b/c puncta in dendrites from Shank3 DD overexpressing neurons (UN) ± tetrodotoxin (TTX) (scale bar = 5 µm). ***B***, Quantification of changes in Shank3+ Homer1b/c+ puncta number induced by scaling up protocol from Shank3 DD overexpressing neurons (number of neurons: DD untreated (UN), n=33, DD TTX, n=24, unpaired t test with Welch’s correction: DD UN vs. DD TTX, ns p=0.6876). ***C***, Quantification of GFP-tagged Shank3 DD intensity changes induced by scaling up protocol from Shank3 DD overexpressing neurons (number of neurons: DD UN, n=33, DD TTX, n=24, unpaired t test with Welch’s correction: DD UN vs. DD TTX, ns p=0.1342). ***D***, Quantification of synaptic Homer1b/c intensity changes induced by scaling up protocol from Shank3 DD overexpressing neurons (number of neurons: DD UN, n=33, DD TTX, n=24, unpaired t test with Welch’s correction: DD UN vs. DD TTX, ns p=0.0664). ***E***, GFP-tagged Shank3 DD intensity plotted against Homer1 intensity for untreated condition. Linear best fit of non-normally distributed raw data shown in dotted line (number of puncta: UN, n=1842). ***F***, GFP-tagged Shank3 DD intensity plotted against Homer1 intensity for 24 hour TTX treatment condition. Linear best fit of non-normally distributed raw data shown in dotted line (number of puncta: TTX, n=1105). ***G***, Log_10_ transformed data from E and F. Line of best-fit for each condition shown in solid lines (F test to determine lines are significantly different, line of best fit slope, ***p<0.0001). Fisher’s z-transformation was used to compare Pearson’s correlation coefficients (***p<0.0001).

**Figure 5.**
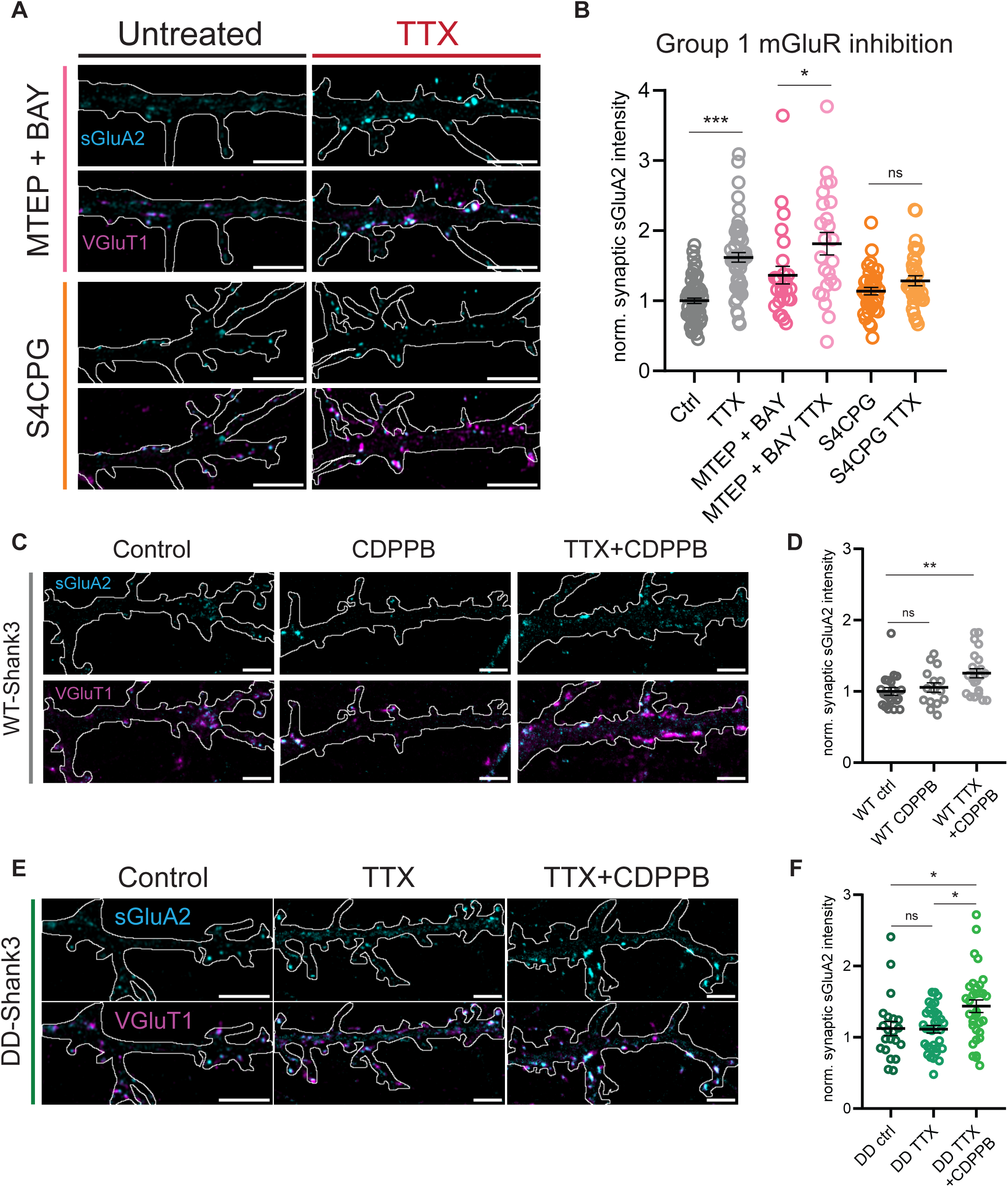
Agonist-dependent group I mGluR signaling is necessary for scaling up and enhanced mGluR5 activity rescues synaptic scaling up in the presence of phosphomimetic Shank3 DD. ***A***, Representative images of synaptic puncta colocalized with sGluA2 and VGLUT1 in neuron dendrites treated with DMSO or 100mM NaOH vehicle (Ctrl) ± tetrodotoxin (TTX), MTEP+BAY ± TTX, and S-4-CPG ± TTX (scale bar = 5 µm) ***B***, Quantification of synaptic sGluA2 intensity changes induced by scaling up protocol (dataset 1: number of neurons: Control, n=29, TTX, n=24, MTEP+BAY, n=26, MTEP+BAY TTX, n=24, Two-way ANOVA with post hoc Tukey’s multiple comparisons test: Ctrl vs. TTX **p=0.0030, Ctrl vs. MTEP+BAY ns p=0.0711, MTEP+BAY vs. MTEP+BAY TTX *p=0.0236; dataset 2: number of neurons: control, n = 32, TTX, n = 30, S-4-CPG, n=36, S-4-CPG TTX, n=33, Two-way ANOVA with post hoc Tukey’s multiple comparisons test: Ctrl vs. TTX ***p<0.0001, Ctrl vs. S-4-CPG ns p=0.5223, S-4-CPG vs. S-4-CPG TTX ns p=0.4529). ***C***, Representative images of synaptic puncta colocalized with sGluA2 and VGLUT1 in dendrites of WT Shank3 overexpressing neurons treated with DMSO vehicle (ctrl) ± CDPPB (CDPPB) ± TTX (CDPPB + TTX) (scale bar = 5 µm). ***D***, Quantification of putative synaptic puncta changes induced by scaling up protocol (number of neurons: WT ctrl, n=21, WT CDPPB, n=15, WT CDPPB + TTX, n=22, Kruskal– Wallis test with post-hoc Dunn’s multiple comparisons test: WT ctrl vs. WT CDPPB, ns p=0.9817, WT ctrl vs. WT CDPPB + TTX, **p=0.0058). ***E***, Representative images of synaptic puncta colocalized with sGluA2 and VGLUT1 in dendrites of DD Shank3 overexpressing neurons treated with DMSO vehicle (ctrl) ± TTX (TTX) ± CDPPB (TTX + CDPPB) (scale bar = 5 µm). ***F***, Quantification of putative synaptic puncta changes induced by scaling up protocol (number of neurons: DD ctrl, n=15, DD TTX, n =31, DD TTX + CDPPB, n=38, Kruskal–Wallis test with post-hoc Dunn’s multiple comparisons test: DD ctrl vs. DD TTX, ns p>0.9999, DD ctrl vs. DD CDPPB + TTX, **p=0.0033, DD TTX vs. DD TTX + CDPPB, *p=0.0115).

### Experimental Design

Experimental conditions were run in parallel with the control condition on sister cultures from the same dissociation. The total puncta intensity measured in experimental conditions was normalized to the mean total puncta intensity of the control in sister cultures, unless otherwise specified.

### Quantification and statistical analysis

All datasets were subjected to a normality test (Shapiro–Wilk test). For data with just a single comparison between two groups (i.e. control vs. TTX), a Mann–Whitney U test was used. For experiments with two or more independent groups (i.e. WT UN vs. P1311L UN, WT UN vs. WT TTX, and P1311L UN vs P1311L TTX), a Kruskall-Wallis test with post-hoc Dunn’s multiple comparisons test was used. For experiments with two independent variables that may have an effect on a dependent variable (i.e. vehicle vs. TTX, vehicle vs. drug alone, drug alone vs. drug+TTX), a two-way ANOVA followed by Tukey’s post hoc correction was used for multiple comparisons. For plots of intensities for colocalized synaptic proteins (figures 1, 2, and 4) which were not normally distributed, Spearman coefficients were used to determine the strength of correlation; the dotted lines on these figures represent the line that fits most of the data. In order to find the statistical line of best fit, the data was log_10_ transformed to achieve normality, and the degree of correlation was found by calculating Pearson’s coefficients. For correlations of log_10_ transformed data, the F-test was used to determine significance between the slopes of best-fit lines for control and TTX conditions. In order to compare correlation coefficients of the log_10_ transformed data, Fisher’s z-transformation was used to determine significant comparisons between control and TTX. Results were considered significant if p < 0.05. Significance symbols used in figures are defined as: *p < 0.05; **p < 0.01; ***p < 0.001; n.s., not significant.

### Data Availability

All data generated in this study are included in the manuscript and available upon request.

## RESULTS

Shank3 is an essential mediator of synaptic scaling, but the signaling pathways downstream of Shank3 phosphorylation that translate a drop in neuronal firing into a homeostatic enhancement of synaptic AMPAR accumulation are largely unknown. Here we ask whether Shank3 and long-form Homer1 interactions, agonist-dependent group I mGluR activity, and downstream group I mGluR canonical signaling pathways are necessary to enable synapses to undergo deprivation-induced synaptic upscaling.

### Synaptic upscaling protocols enhance excitatory synaptic density, and produce correlated changes in pre- and post-synaptic proteins

Previous work has shown that synaptic scaling is accompanied by changes in the abundance of many postsynaptic proteins, including Shank3 and Homer1 (Desch et al., 2021; Heavner et al., 2021; Wu et al., 2022; Yuan et al., 2023). In addition to modulating postsynaptic strength, some studies have also reported changes in synapse number and presynaptic protein accumulation (Glebov et al., 2017; Dörrbaum et al., 2020), but these effects are often smaller and more variable than changes in AMPAR accumulation. Here we compiled a large dataset (control n=120, TTX treated n =104 neurons) to determine the degree to which these pre- and postsynaptic changes are coupled. We induced synaptic upscaling by treating rat visual cortical cultures (used for all subsequent experiments, unless otherwise stated) with TTX for 24 hours to suppress firing, transfected neurons at low efficiency with a fluorescent protein to allow us to carefully quantify synaptic contacts onto individual pyramidal neurons, then fixed and labeled them using antibodies directed against a surface epitope of the postsynaptic GluA2 subunit of the AMPAR (sGluA2) and the presynaptic glutamate transporter (VGluT1, Fig. 1A). The density of colocalized sGluA2 and VGluT1 puncta, which represent putative synaptic contacts, as well as the intensity of the fluorescent signals at these colocalized sites, was then quantified along apical-like pyramidal neuron dendrites. The postsynaptic accumulation of sGluA2 in TTX-treated neurons was enhanced compared to vehicle-treated neurons from sister cultures (Fig. 1A,B), as expected (Ibata et al., 2008; Tatavarty et al., 2013; Gainey et al., 2015; Wu et al., 2022). Further, there was a small but significant increase in synaptic density (Fig. 1C), and a small but significant increase in the intensity of VGluT1 at presynaptic sites (Fig. 1D). To determine whether TTX produces correlated changes in pre- and postsynaptic markers at individual synapses, we plotted the intensity of VGluT1 vs sGluA2 for each synaptic contact for the control (Fig.1E, n = 4398) and TTX conditions (Fig. 1F, n = 4658). Under both conditions there was a strong correlation (control r^2^ = 0.801 vs. TTX r^2^ = 0.785); however, the line-of-best-fit slope of the relationship in the TTX condition is significantly steeper than control (Fig. 1G, Control slope = 0.883 vs TTX slope = 0.927), consistent with a larger postsynaptic effect of synaptic scaling on sGluA2 accumulation than on VGlut1 levels (compare 1B and 1D). Thus, synaptic upscaling protocols induce coupled pre- and postsynaptic changes, as well as an increase in synapse number.

### Synaptic upscaling protocols enhance endogenous Shank3-Homer1 colocalization

Shank3 and Homer1 are known binding partners (Cheng et al., 2006) and both have been implicated in synaptic upscaling (Clifton et al., 2019; Tatavarty et al., 2020; Bockaert et al., 2021; Heavner et al., 2021; Wu et al., 2022). We therefore wished to explore how TTX treatment affects their colocalization at synaptic sites. To this end, we performed immunocytochemistry against Shank3 and Homer1b/c (using an antibody directed against a sequence absent in Homer1a), and quantified the signal at colocalized sites within the apical dendrites of GFP-expressing neurons. The density of colocalized Shank3 and Homer1b/c puncta was approximately doubled after TTX treatment (Fig. 2A,B). Additionally, both Shank3 and Homer1b/c intensities at colocalized puncta were significantly increased in the TTX condition to a similar degree (Fig. 2C,D). Finally, we examined the correlation between Shank3 and Homer1b/c intensity at individual puncta from control (Fig. 2E, n = 1511) and TTX treated neurons (Fig. 2F, n = 2228). For both conditions, Shank3 and long-form Homer1 intensities were significantly correlated with each other, and the slopes of both relationships were similar and significantly different from zero (Fig. 2E,F). Interestingly, synaptic scaling was accompanied by a significant increase in both the slope of this relationship and the strength of correlation, suggesting that synaptic upscaling is accompanied by a greater and more tightly regulated association between Shank3 and long-form Homer1. Thus, while the short-form Homer1a has been linked to synaptic downscaling (Hu et al., 2010), these data implicate long-form Homer1 in synaptic upscaling.

### Phosphomimetic Shank3 DD has reduced interaction with Homer1b/c but maintains its ability to localize to the synapse

Synaptic scaling reduces Shank3 phosphorylation at S1586/S1615, and overriding this dephosphorylation by overexpressing the phosphomimetic mutant Shank3 DD (Fig. 3A) blocks upscaling; in contrast, scaling up is preserved after overexpression of the phosphodeficient mutant Shank3 AA (Wu et al., 2022). We thus wondered whether the phosphorylation state of Shank3 might influence its ability to directly bind long-form Homer1. To test this, we co-transfected HEK-T cells with Homer1c plus one of several forms of GFP-tagged Shank3 (Fig. 3A): wildtype (WT); phosphodeficient (AA); phosphomimetic (DD); or P1311L, a point mutant known to have significantly reduced binding to Homer1 proteins (Tu et al., 1999). Dynabeads bound to GFP antibody or GFP-trap magnetic beads/particles were used to pull down Shank3 protein from cell lysates, and Western blots were then performed to quantify the ratio of Homer1c to Shank3 (Fig. 3B). Shank3 WT and AA were able to pull down Homer1c with similar efficacy (AA was 88% ± 3.8% of WT) while the negative control P1311L had a greatly reduced efficacy (P1311L was 25.5% ± 6.5% of WT). Intriguingly, the phosphomimetic Shank3 DD mutant also had reduced efficacy in pulling down Homer1c (DD was 61.7% ± 4.8% of WT, Fig. 3B,C).

Next we assessed the ability of Shank3 DD and Shank3 P1311L to interfere with upscaling. Consistent with previous work (Wu et al., 2022), neurons expressing Shank3 DD were unable to express upscaling; in contrast, neurons overexpressing the point mutant Shank3 P1311L, exhibited normal synaptic scaling (Fig. 3D,E). We next examined the synaptic localization of both forms of mutant Shank3 relative to WT, by examining the fraction of putative synaptic puncta (defined as sites of colocalization of sGluA2 and VGluT1) that contain tagged Shank3. Interestingly, while Shank3 DD localized to synaptic sites to the same degree as WT Shank3, Shank3 P1311L had greatly reduced synaptic localization (Fig. 3F,G). This reduced synaptic localization may explain why Shank3 P1311L is not able to prevent synaptic scaling up, as this effect presumably relies on displacing endogenous Shank3 from synapses (Fig. 3E). Finally, we verified that overexpressing these Shank3 mutants did not affect the density of puncta colocalizing sGluA2 and VGluT1; indeed we found no significant differences in the density of putative synapses between neurons overexpressing Shank3 WT, DD, and P1311L (Fig. 3H).

### Synaptic upscaling protocols do not alter Shank3-Homer1 colocalization or produce correlated changes between these proteins in the presence of phosphomimetic Shank3 DD

If Shank3 phosphorylation at S1586/S1615 disrupts the association between Shank3 and long-form Homer1 at synapses, we would expect that TTX treatment would no longer enhance Homer1 accumulation in the presence of Shank3 DD. To test this, we transfected sister cultures at low efficiency with GFP-tagged Shank3 DD, and quantified the degree of colocalization of total Shank3 (endogenous and overexpressed) and endogenous Homer1b/c. In the presence of Shank3 DD, TTX treatment did not significantly increase the density of colocalized puncta (Fig. 4A,B). Further, the intensity of Shank3 and Homer1b/c signals at colocalized sites was not altered by TTX treatment (Fig. 4C,D).

Next, we examined the correlation between Shank3 and Homer1b/c intensity at individual puncta from neurons expressing Shank3 DD, without (Fig. 4E, n = 1842) or with (Fig. 4F, n = 1105) TTX treatment. In the untreated condition, Shank3 and long-form Homer1 intensities were significantly correlated with each other (Fig. 4E). Strikingly, while this correlation normally increases after TTX treatment (Fig. 2E, F), in the presence of Shank3 DD there was instead a robust decrease in correlation after TTX treatment (Fig.4E,F: from r2 = 0.881 to r2 = 0.779), as well as a significant decrease in slope (Fig.4G: from 0.322 to 0.247). Thus, Shank3 DD disrupts the ability of activity-blockade to enhance the association between Shank3 and long-form Homer1. Together, these data suggest that Shank3 DD may block synaptic upscaling by preventing the enhanced association between Shank3 and long-form Homer1.

### Agonist-dependent group I mGluR signaling is necessary for scaling up and enhanced mGluR5 activity rescues synaptic scaling up in the presence of phosphomimetic Shank3 DD

Long-form Homer1 links Shank proteins, including Shank3, to group I mGluRs to modulate signaling (Wang et al., 2016b). We thus wondered whether Shank3 DD might block scaling up by preventing activation of group I mGluRs. Group I mGluRs are present at the perisynapse and have both agonist-dependent and agonist-independent activity (Ango et al., 2001; Bockaert et al., 2021; Mango and Ledonne, 2023). It is known that group I mGluR noncompetitive inhibitors, specifically those targeting agonist-independent signaling, can abolish homeostatic synaptic downscaling (Hu et al., 2010), but a role for mGluR signaling in upscaling has not been assessed. Therefore, we began by testing the ability of the non-competitive inhibitors BAY 36-7620 (mGluR1) and MTEP (mGluR5) to block synaptic upscaling. Co-treatment of cultures with TTX and these noncompetitive inhibitors for 24 hours resulted in an increase in synaptic AMPAR accumulation of similar magnitude to that produced by TTX alone (Fig. 5A,B: MTEP+BAY vs MTEP+BAY TTX), indicating that agonist-independent activity of group I mGluRs is not necessary for synaptic upscaling to occur.

Agonist-independent activity is important for downscaling (Hu et al., 2010), and can result from binding to short-form Homer1a (Ango et al., 2001; Clifton et al., 2019). We next wondered whether, in contrast, agonist-*dependent* activity of the group I mGluRs might be the critical signal needed for synaptic upscaling. To test this, we co-treated cultures for 24 hours with TTX and S-4-CPG, a specific competitive inhibitor of group I mGluRs. We found that competitive inhibition led to a complete loss of synaptic upscaling (Fig. 5A,B: S4CPG vs S4CPG TTX). Thus, agonist-dependent, rather than agonist-independent, activity of group I mGluRs is necessary for homeostatic synaptic upscaling.

We showed previously that Shank3 DD traffics correctly to synaptic sites and blocks upscaling (Wu et al., 2022); if this effect is mediated through a reduction in group I mGluR signaling, then enhancing this signaling should rescue scaling up in the presence of Shank3 DD. Because the two group I mGluRs, mGluR1 and mGluR5, have some distinct and some overlapping functions and actions, we decided to selectively target mGluR5 with CDPPB, a positive allosteric modulator of mGluR5. CDDPB binds to a site on mGluR5 that is distinct from the ligand binding site, to enhance the response to binding of the endogenous ligand, glutamate. We first ensured that CDPPB does not affect upscaling in the presence of WT Shank3 overexpression; in neurons expressing WT Shank3, synaptic sGluA2 intensity was unaffected by CDPPB at baseline, and TTX induced the normal increase in the presence of CDPPB (Fig. 5C,D). When we overexpressed Shank3 DD and treated neurons with TTX, synaptic upscaling was absent (Fig. 5E,F), as expected (Wu et al., 2022). However, co-treatment of Shank3 DD overexpressing neurons with CDPPB and TTX induced robust synaptic upscaling (Fig. 5E,F). Thus enhancement of mGluR5 function is sufficient to rescue synaptic scaling up; further, mGluR5 signaling is downstream of Shank3 phosphorylation in the signaling pathways mediating synaptic scaling up.

### Canonical group I mGluR downstream signaling pathways are necessary for synaptic scaling up

We next wanted to investigate which pathway(s) downstream of group I mGluRs are necessary for homeostatic synaptic upscaling. The canonical pathway(s) downstream of group I mGluRs is driven by Gq/G11-dependent activation of PLC that leads to PKC activation and also to mobilization of intracellular calcium from internal stores (Conn and Pin, 1997; Hermans and Challiss, 2001; Mango and Ledonne, 2023). Here we sought to interrupt several steps in this canonical pathway, focusing on those for which selective inhibitors are available, to determine their necessity during synaptic upscaling.

To target PLC, we used the selective PLC inhibitor, aminosteroid U73122 (Kamato et al., 2015; Neacsu et al., 2020). We found that TTX was unable to increase synaptic AMPAR accumulation in the presence of U73122 (Fig. 6A,B, U73122 vs U73122 + TTX), indicating that synaptic scaling is impaired under these conditions. One target of PLC activation is Ca^2+^ release from intracellular stores, which is modulated by the sarcoplasmic-endoplasmic reticulum Ca^2+^ ATPase (SERCA-ATPase) (Masu et al., 1991; Abe et al., 1992; Putney and Tomita, 2012). We used the selective SERCA ATPase inhibitor cyclopiazonic acid (CPA), which has been shown to deplete internal calcium stores in hippocampal neurons (Eder and Bading, 2007), to target this pathway. Because it takes time to deplete intracellular stores of calcium, the SERCA-ATPase inhibitor CPA was added to the culture media 6 hours before TTX, and cultures were fixed and labeled 24 hours after TTX addition. Interestingly, there was a significant increase in the sGluA2 synaptic intensity for the CPA+TTX condition compared to the CPA only condition (Fig. 6A,B); suggesting that intact internal calcium stores are not essential for synaptic upscaling.

**Figure 6.**
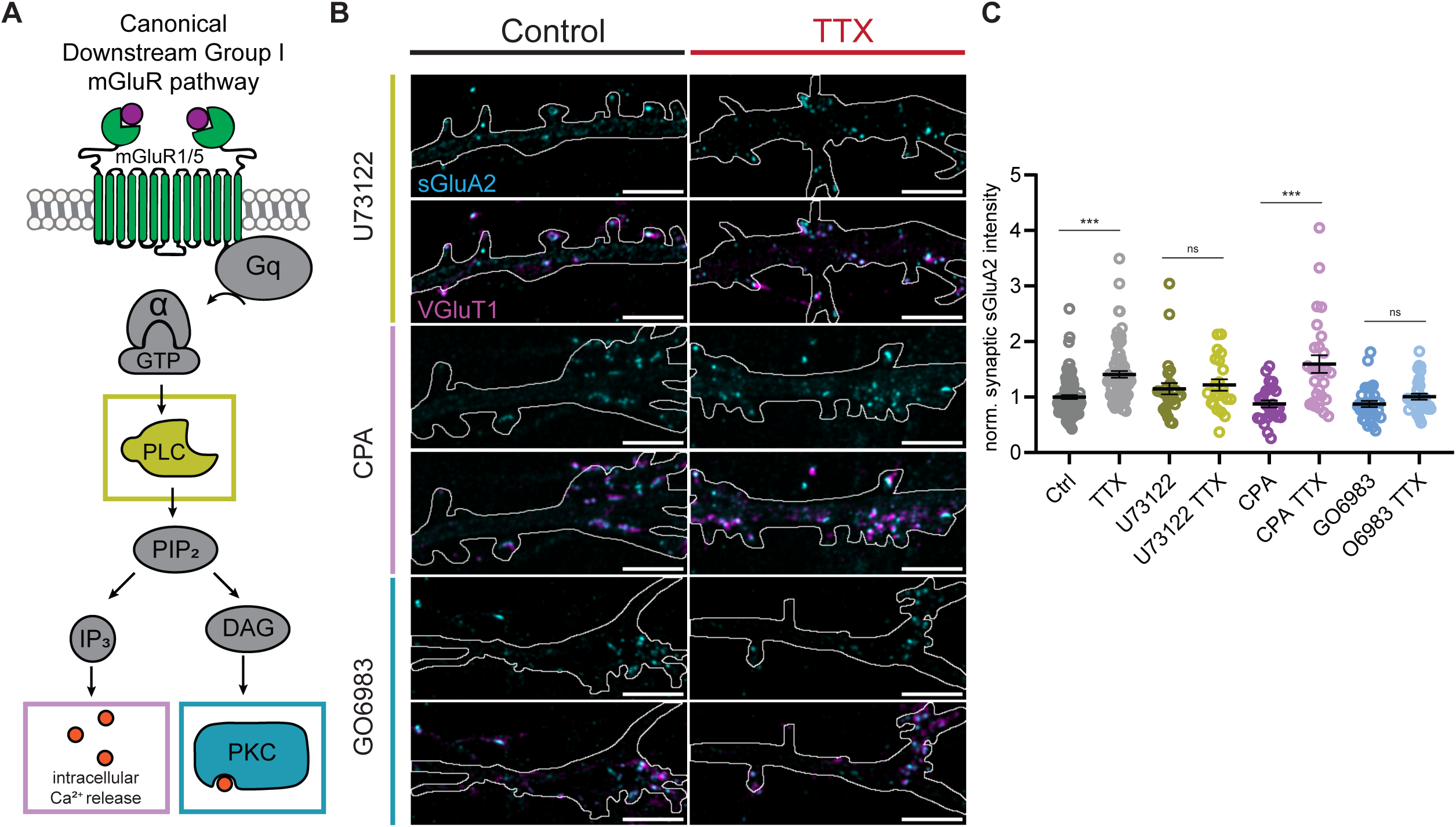
Canonical group I mGluR downstream signaling pathways are necessary for synaptic scaling up. ***A***. Representative images of synaptic puncta colocalized with sGluA2 and VGLUT1 in neuron dendrites treated with DMSO vehicle (Ctrl) ± tetrodotoxin (TTX), and U73122 ± TTX, CPA ± TTX, and GO6983 ± TTX (scale bar = 5 µm). ***B***, Quantification of synaptic sGluA2 intensity changes induced by scaling up protocol (dataset 1: number of neurons: control, n=26, TTX, n=22, U73122, n=27, U73122 TTX, n=23, Two-way ANOVA with post hoc Tukey’s multiple comparisons test: Ctrl vs. TTX **p=0.0093, Ctrl vs. U73122 ns p=0.6709, U73122 vs. U73122 TTX ns p=0.9592; dataset 2: number of neurons: control, n = 35, TTX, n=29, CPA, n=26, CPA TTX, n=27, Two-way ANOVA with post hoc Tukey’s multiple comparisons test: Ctrl vs. TTX *p=0.0253, Ctrl vs. CPA ns p=0.8416, CPA vs. CPA TTX ***p<0.0001; dataset 3: number of neurons: control, n = 30, TTX, n = 25, GO6983, n=31, GO6983 TTX, n=33, Two-way ANOVA with post hoc Tukey’s multiple comparisons test: Ctrl vs. TTX ***p=0.0001, Ctrl vs. GO6983ns p=0.3848, GO6983 vs. GO6983 TTX ns p=0.3265).

Finally, we targeted the other major target of the Gq/G11-mediated group I mGluR signaling, PKC. To test for a role of PKC, we used the broad-spectrum PKC inhibitor, GO6983, to target all isoforms of PKC (Gschwendt et al., 1996; Zheng et al., 2000). In the presence of GO6983, TTX was unable to increase synaptic sGluA2 relative to inhibitor GO6983 alone (Fig. 6A,B). Taken together, these findings show that multiple branches of the group I mGluR canonical pathway signaling cascade are necessary for the induction of synaptic scaling up.

## DISCUSSION

While synaptic scaling has been studied extensively in a wide variety of contexts (Desai et al., 2002; Wierenga et al., 2005; Goel and Lee, 2007; Ibata et al., 2008; Steinmetz and Turrigiano, 2010; Sun and Turrigiano, 2011; Tatavarty et al., 2013; Gainey et al., 2015; Steinmetz et al., 2016; Wu et al., 2022), the molecular pathways necessary to induce the complex synaptic reorganization and changes in synaptic AMPAR accumulation that drive it are still being defined. In particular, while Shank3 dephosphorylation is necessary for upscaling (Wu et al., 2022), the pathways downstream of Shank3 dephosphorylation are unknown. Here, we utilize genetic and pharmacological manipulations to show that agonist-dependent group I mGluR activity, and key components of its downstream canonical signaling pathways, are required for upscaling. Importantly, the Shank3 phosphorylation state influences binding interactions with long-form Homer1, suggesting that the impact of Shank3 phosphorylation state on mGluR activity is achieved through the modulation of Shank3/long-form Homer1 interactions. Together, our findings support a unified model in which the same molecular players known to be crucial for synaptic downscaling during activity enhancement – Shank3, Homer1 isoforms, and group I mGluRs – are reconfigured by activity deprivation to enable the distinct and complementary signaling required for synaptic upscaling.

Previous work has shown that synaptic scaling down relies on agonist-independent group I mGluR activity (Hu et al., 2010; Diering et al., 2017). In this model of downscaling, high network activity leads to an increase in the immediate early gene Homer1a; Homer1a then functions as a dominant negative to prevent the crosslinking of synaptic proteins, including Shank3 and group I mGluRs. By displacing long-form Homer1, Homer1a is able to bind group I mGluRs to evoke mGluR constitutive, or agonist-independent, activity. This constitutive mGluR activity is required for synaptic downscaling, through a signaling cascade that involves phosphorylation of extracellular signal–regulated kinase 1/2 (ERK1/2, Hu et al., 2010; Diering et al., 2017).

Here, we define a complementary signaling pathway during synaptic upscaling, that uses activity-dependent changes in the phosphoproteome to rearrange the same signaling elements from a configuration that enables downscaling into one that instead enables upscaling. We find that prolonged activity deprivation enhances the association between Shank3 and long-form Homer1, in a manner that depends on the Shank3 phosphorylation state. Moreover, we find that competitive, rather than noncompetitive, inhibition of group I mGluRs prevents the expression of synaptic upscaling. Competitive inhibition targets both agonist-dependent and -independent signaling, while noncompetitive inhibition only targets agonist-independent signaling. Therefore, our results suggest that agonist-dependent activity is necessary for synaptic upscaling. Together with previous work, our data suggest that up- and down-scaling protocols result in several reciprocal changes that influence Shank3/Homer1/mGluR associations: first, they alter the Shank3 phosphorylation state and Homer1 short-form abundance to bias Shank3/Homer associations toward Homer1 long-form (upscaling) or short-form (downscaling); and second, as a consequence of these altered associations, mGluR5 signaling is switched from agonist dependent (upscaling) to agonist independent (downscaling) modes.

Group I mGluR activation can initiate signaling through a number of downstream effectors. Because Homer1a displaces group I mGluRs from the major signaling effectors inositol 1,4,5-triphosphate and protein kinase C (IP3R/PKC, Brakeman et al., 1997), this likely biases signaling towards other pathways such as ERK1/2, that have been implicated in synaptic downscaling (Diering et al., 2017). Binding of long-form Homer and group I mGluRs results in a switch from constitutive to agonist-dependent mGluR activity (Fagni et al., 2003), consistent with our observation that upscaling requires agonist-dependent mGluR signaling and activation of downstream PLC and PKC signaling. Interestingly, although intracellular calcium release regulated by SERCA-ATPase, IP3-receptors, and Ryanodine receptors is important in mGluR-dependent forms of LTP and LTD (Holbro et al., 2009; Lüscher and Huber, 2010; Wang et al., 2016a; Chami and Checler, 2020), it is not necessary for synaptic upscaling, indicating that distinct forms of synaptic plasticity utilize different downstream mGluR signaling pathways.

There are a host of molecular players that have been implicated in synaptic scaling up (Pozo and Goda, 2010; Wen and Turrigiano, 2024), suggesting that Shank3/Homer1/mGluR5 signaling likely works in concert with additional signaling and trafficking pathways to enhance synaptic AMPAR accumulation. Since mGluR signaling is able to regulate transcription, one possibility is that this pathway could be important for transcriptional changes necessary for synaptic upscaling (Ibata et al., 2008; Steinmetz et al., 2016; Schaukowitch et al., 2017). Another interesting possibility is that Shank3/Homer1/mGluR complexes might influence Shank3 binding to GRIP1 (Sheng and Kim, 2000), to enable GRIP1 to capture AMPARs at the PSD (Gainey et al., 2015; Tan et al., 2015). An important point is that mGluR5 signaling is not sufficient to induce scaling up; rather, this pathway, like upstream Shank3 phosphorylation, serves to gate the direction of homeostatic compensation.

As a key part of the tripartite synapse, glia perform crucial functions that support synaptic plasticity in neurons (Ben Achour and Pascual, 2010; Panatier and Robitaille, 2016; Sancho et al., 2021), and group I mGluRs are present in glial cells (Muly et al., 2003). One of the most well-characterized mechanisms of astrocytic intracellular Ca2+ signaling is the canonical PLC and IP3 pathway; when GPCRs such as mGluRs are activated, the coupled Gq protein leads to IP3 receptor activation and calcium release from the endoplasmic reticulum (ER, Agulhon et al., 2008; Volterra et al., 2014; Bazargani and Attwell, 2016; Shigetomi et al., 2016). However, astrocytic calcium signaling has been described in the cytosol of the soma in response to neuronal activity (Wang et al., 2006; Winship et al., 2007; Schipke et al., 2008; Ding et al., 2013; Lind et al., 2013). Furthermore, recent studies have also established calcium signaling in astrocytic endfeet and synapse-associated processes in response to neuronal activity (Shigetomi et al., 2013; Otsu et al., 2015; Agarwal et al., 2017; Bindocci et al., 2017; Stobart et al., 2018). Thus, glia are responsive to increases in neuronal activity, but it is not clear whether activity blockade also influences neuron-glial signaling. At the moment, our data cannot rule out a role for astrocytic mGluR signaling in synaptic scaling, but past data finds ERK1/2 phosphorylation in synaptosomal fractions is specifically increased (Diering et al., 2017), suggesting that mGluR signaling cascades mediating downscaling are localized within neurons. Since we show that altered Shank3 phosphorylation and interactions with Homer1 occur within neurons, the most likely scenario is that mGluR signaling during scaling induction is also occurring within neurons.

Our data show that TTX-induced synaptic scaling up requires agonist-dependent mGluR signaling. This raises the interesting question of what drives agonist-dependent activity of group I mGluRs in the presence of TTX, which blocks spike-mediated glutamate release. One likely possibility is that spike-independent, spontaneous vesicular release of glutamate is sufficient to activate mGluR signaling during activity-blockade. Given the predominantly perisynaptic localization of group I mGluRs (Scheefhals and MacGillavry, 2018; Scheefhals et al., 2023), and reports that spontaneous release happens in areas that encompass synaptic and perisynaptic sites (Tang et al., 2016; Farsi et al., 2021), spontaneous release may be well-placed to activate these receptors. Taken together, our work suggests a model in which activity-dependent changes in Shank3 phosphorylation modulate Shank3 interactions with long-form Homer1, leading to a switch from constitutive to agonist-dependent activity of group I mGluRs to enable synaptic scaling up. This supports the possibility that ASD phenotypes arising from *Shank3* loss (Moessner et al., 2007; Zhou et al., 2016; Bey et al., 2018) may be caused in part through disrupted signaling through group I mGluRs, as has been suggested for other ASD-associated genes such as FMR1 (Huber et al., 2002; Nakamoto et al., 2007; Iliff et al, 2013).

**Table 1.** Two-way ANOVA for MTEP+BAY & S-4-CPG experiments.

| ANOVA Table | SS (Type III) | Degrees of Freedom | Mean Squares | F (DFn, DFd) | P value |
| --- | --- | --- | --- | --- | --- |
| **Interaction (MTEP+BAY)** | 0.05378 | 1 | 0.05378 | F (1, 99) = 0.1793 | P=0.6729 |
| **Effect of TTX on Veh** | 6.273 | 1 | 6.273 | F (1, 99) = 20.92 | P<0.0001 |
| **Effect of TTX on MTEP+BAY** | 2.611 | 1 | 2.611 | F (1, 99) = 8.708 | P=0.0040 |
| **Residual (MTEP+BAY)** | 29.69 | 99 | 0.2999 |  |  |
| **Interaction (S-4-CPG)** | 2.324 | 1 | 2.324 | F (1, 127) = 13.41 | P=0.0004 |
| **Effect of TTX on Veh** | 5.625 | 1 | 5.625 | F (1, 127) = 32.47 | P<0.0001 |
| **Effect of TTX on S-4-CPG** | 0.5401 | 1 | 0.5401 | F (1, 127) = 3.117 | P=0.0799 |
| **Residual (S-4-CPG)** | 22.00 | 127 | 0.1732 |  |  |

**Table 2.** Two-way ANOVA for U73122, CPA, and GO6983 experiments.

| **ANOVA Table** | **SS (Type III)** | **Degrees of Freedom** | **Mean Squares** | **F (DFn, DFd)** | **P value** |
| --- | --- | --- | --- | --- | --- |
| **Interaction (U73122)** | 0.8607 | 1 | 0.8607 | F (1, 93) = 3.754 | P=0.0557 |
| **Effect of TTX on Veh** | 1.597 | 1 | 1.597 | F (1, 93) = 6.968 | P=0.0097 |
| **Effect of TTX on U73122** | 0.03907 | 1 | 0.03907 | F (1, 93) = 0.1704 | P=0.6807 |
| **Residual (U73122)** | 21.32 | 93 | 0.2293 |  |  |
| **Interaction (CPA)** | 1.145 | 1 | 1.145 | F (1, 117) = 3.466 | P=0.0651 |
| **Effect of TTX on Veh** | 8.027 | 1 | 8.027 | F (1, 117) = 24.30 | P<0.0001 |
| **Effect of TTX on CPA** | 0.1629 | 1 | 0.1629 | F(1, 117) = 0.4932 | P=0.4839 |
| **Residual (CPA)** | 38.64 | 117 | 0.3303 |  |  |
| **Interaction (G06983)** | 0.4209 | 1 | 0.4209 | F (1, 115) = 4.427 | P=0.0376 |
| **Effect of TTX on Veh** | 1.854 | 1 | 1.854 | F (1, 115) = 19.50 | P<0.0001 |
| **Effect of TTX on G06983** | 1.776 | 1 | 1.776 | F (1, 115) = 18.68 | P<0.0001 |
| **Residual (G06983)** | 10.93 | 115 | 0.09507 |  |  |

## ACKNOWLEDGEMENTS

This work was supported by a Howard Hughes Medical Institute Gilliam Fellowship (AAG) and NIH R35NS111562 (GGT). We thank the Brandeis Light Microscopy Core Facility RRID:SCR_025892 for equipment (Zeiss LSM880 and LSM880 Fast AiryScan confocal microscopes) and technical assistance. We thank Chi-Hong Wu for invaluable mentorship, Lauren Tereshko for the Metamorph colocalization analysis scripts, and Lirong Wang for her work generating neuronal cultures.

---

**Supplemental Figure 1.**
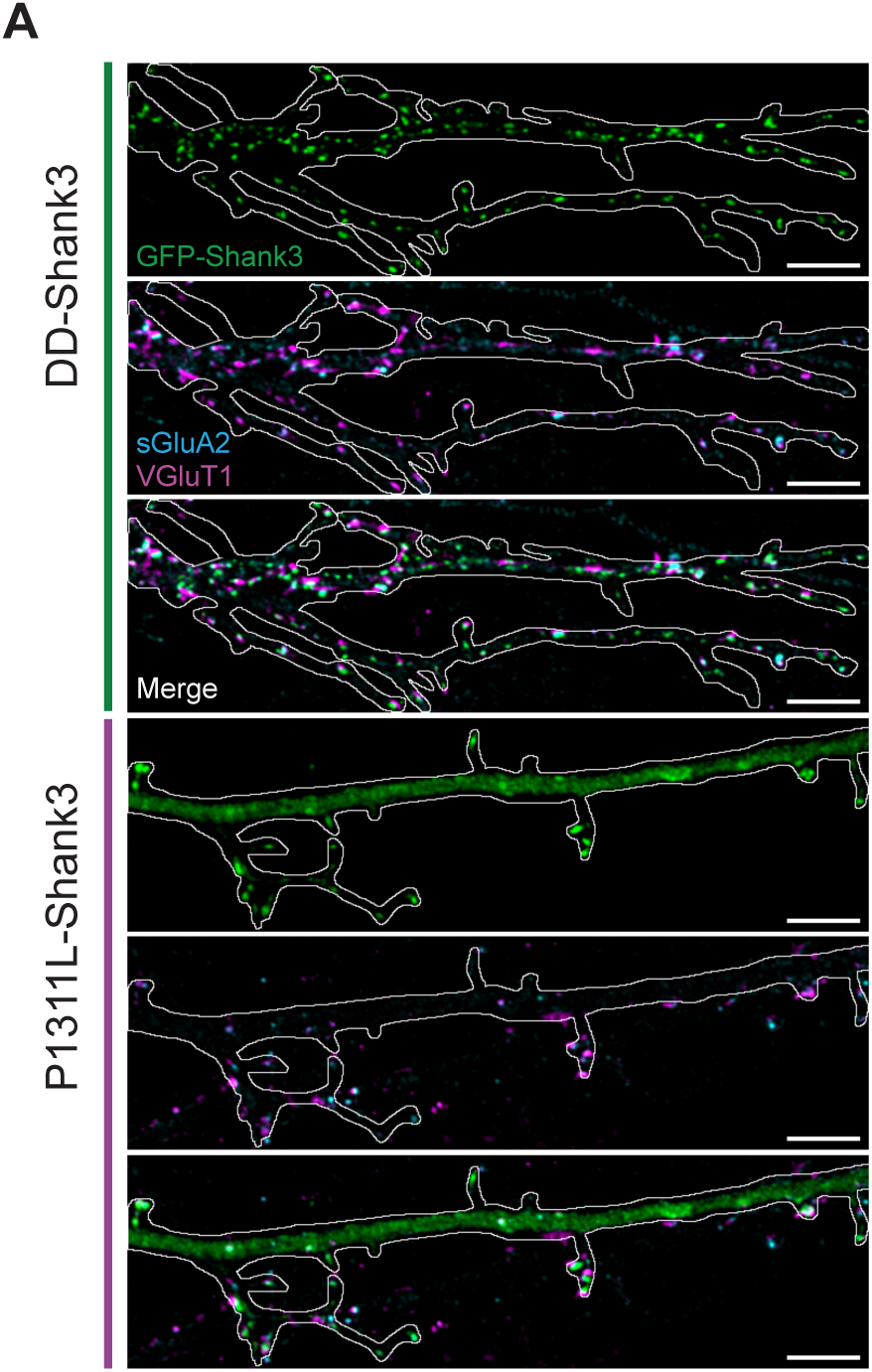
Overexpression of Shank3 P1311L causes decreased synaptic localization of Shank3 at baseline compared to Shank3 DD and Shank3 WT. ***A***.Representative images of Shank3 P1311L and Shank3 DD and synaptic puncta colocalized with sGluA2 and VGLUT1 in neuron dendrites at baseline (scale bar = 5 µm).

